# Apramycin efficacy in the treatment of carbapenem-resistant *Enterobacterales* in murine blood stream infection models

**DOI:** 10.1101/2023.12.10.570991

**Authors:** Niels Frimodt-Møller, Jon U. Hansen, Michel Plattner, Douglas L. Huseby, Stine Radmer Almind, Klara Haldimann, Marina Gysin, Anna Petersson, Onur Ercan, Lea Ganz, Diarmaid Hughes, Carina Vingsbo Lundberg, Sven N. Hobbie

**Author notes:** these authors share senior authorship (Niels Frimodt-Møller, Diarmaid Hughes, Carina Vingsbo Lundberg, Sven N. Hobbie).

## Abstract

**Background:** The aminoglycoside apramycin has been proposed as a drug candidate for the treatment of critical Gram-negative systemic infections. However, its potential in the treatment of drug-resistant bloodstream infections (BSIs) has yet to be assessed.

**Methods:** The resistance gene annotations of 26 493 blood culture isolates were analyzed. *In vitro* profiling of apramycin comprised cell-free translation assays, broth microdilution, and frequency of resistance determination. The efficacy of apramycin was studied in a mouse peritonitis model for nine *E. coli* and *K. pneumoniae* isolates.

**Results:** Genotypic aminoglycoside resistance was identified in 87.8% of all 6973 carbapenem-resistant *Enterobacterales* blood-culture isolates, in comparison to 46.4% of colistin and 2.1% of apramycin resistance. Apramycin activity against methylated ribosomes was > 100-fold higher than other aminoglycosides. Frequencies of resistance were < 10^−9^ at 8 × MIC. Tentative epidemiological cutoffs (ECOFFs) were determined as 8 μg/mL for *E. coli* and 4 μg/mL for *K. pneumoniae*. A single dose of 5 to 13 mg/kg resulted in a 1-log CFU reduction in the blood and peritoneum. Two doses of 80 mg/kg, resulting in an exposure that resembles the AUC observed for a single 30 mg/kg dose in humans, resulted in complete eradication of carbapenem- and aminoglycoside-resistant bacteremias.

**Conclusion:** Encouraging coverage and potent in-vivo efficacy against a selection of highly drug-resistant *Enterobacterales* isolates in the mouse peritonitis model warrants further consideration of apramycin as a drug candidate for the treatment and prophylaxis of BSI.

## 1. Introduction

Aminoglycoside antibiotics play an important role in the treatment of serious systemic infections such as blood stream infections (BSI), endocarditis, pneumonia, pyelonephritis, and other complicated urinary tract infections [1]. In parallel to the increase in Gram-negative pathogens of resistance towards β-lactam antibiotics (production of extended spectrum β-lactamases (ESBLs and carbapenemases)), fluoroquinolones, trimethoprim and sulfonamides, there has also been an increase in resistance towards standard aminoglycosides such as gentamicin, tobramycin, and amikacin; mostly based on a spread of genes encoding aminoglycoside-modifying enzymes [2]. The use of aminoglycosides is hampered by nephro- and ototoxicity, which are preceded by drug accumulation in the kidneys and the cochlea. Since this accumulation is indicated by increasing drug trough plasma concentrations therapeutic drug monitoring is considered necessary especially with treatment extending beyond three days [1]. This increases the workload and cost of treatment with these antibiotics.

Apramycin is an aminoglycoside discovered as nebramycin factor 2 and has since almost exclusively been used in animal husbandry [3]. Due to recent findings of the stability of apramycin against most aminoglycoside-resistance transferases as well as other mechanisms found in clinical isolates world-wide, pharmaceutical-grade apramycin (EBL-1003) has been proposed as a next-generation aminoglycoside for human clinical use, active against so-called pan-aminoglycoside resistant isolates [4-8]. Notably, EBL-1003 has displayed lower nephro- and ototoxicity than other aminoglycosides [4, 9, 10], and has demonstrated encouraging efficacy in the thigh, lung, and UTI animal infection models [10-15]. However, the efficacy of EBL-1003 against BSI in animal infection models has yet to be studied.

The purpose of this study was to expand on studying the activity of apramycin against blood stream infections caused by carbapenemase-producing *Enterobacterales* (CPE). The methods included in-silico data base analysis for genotypic susceptibility, in-vitro translation inhibition assays, in-vitro mutational resistance analysis and broth microdilution studies with wild-type and carbapenemase-producing multi-drug resistant *E. coli* and *K. pneumoniae* clinical isolates. Further, to elucidate the efficacy of EBL-1003 in the treatment of BSIs caused by *E. coli* or *K. pneumoniae* it was tested in the mouse sepsis/peritonitis model, for a total of nine MDR isolates causing BSI in the mouse sepsis/peritonitis model.

## 2. Material and methods

### 2.1. Data base analysis for aminoglycoside resistance genes in blood culture isolates

An epidemiologic data base analysis for genotypic susceptibility of blood culture isolates was done as described previously for respiratory and uropathogens [10, 14]. Data was downloaded from the NCBI National Database of Antibiotic Resistant Organisms (NDARO) on May 02, 2023 (https://www.ncbi.nlm.nih.gov/pathogens/antimicrobial-resistance) [16]. The inclusion criteria were “host: homo sapiens”, “isolation type: clinical” and “isolation source: blood, blood culture, human blood, human blood culture, bloodstream, bloodstream isolates and sepsis”. Resistance gene annotations for 40 888 genomes not including *Salmonella enterica* were analyzed for the presence of known antibiotic resistance determinants for aminoglycosides, colistin, and carbapenems (**Table S1**). The search included variants, subvariants and mutants of all resistance gene families. Genotypic susceptibility was inferred from the presence or absence of specific resistance determinants.

### 2.2. Bacterial strains

Wild type *E. coli* (*n* = 115) and *K. pneumoniae* (*n* = 125) clinical isolates were sourced at four different study sites: Uppsala University (*n* = 73), the University of Zurich (*n* = 69), the Rigshospitalet in Copenhagen (*n* = 58), and IHMA Europe (Monthey, Switzerland; *n* = 40) (**Table S2**). Fifteen *Enterobacterales* isolates with high-level resistance to aminoglycoside antibiotics in clinical use were provided by the sites in Uppsala and Zurich (**Table S3**). In addition, twenty *Enterobacterales* isolates representative of a diversity of carbapenemase genotypes were provided by the Statens Serum Institute (**Table S4**) [17]. *E. coli* ATCC 25922 was used as quality control and *E. coli* reference strain. *K. pneumoniae* strains ATCC 13883 and SSI 3010 were used as *K. pneumoniae* wild-type reference strains.

### 2.3. In-vitro ribosomal translation inhibition assays

For the ribosomal inhibition assays, *E coli rmtB* isolate EN0591 and *K. pneumoniae rmtB* isolate EN0593 (**Table S3**) were used in the preparation of S30 extract for *in vitro* translation inhibition assays as described previously [18]. In brief, m^7^G1405-methylated ribosomes were extracted from bacterial cells with a microfluidizer processor (Microfluidics, Westwood, MA, USA) at 25 000 lb/in^2^, addition of dithiothreitol (DTT) to 1 mM, and centrifugation at 30 000 × *g* at 4°C. Translation reaction mixtures containing 4 μL of the S30 extract, 0.2 mM amino acid mix, 6 μg tRNA (Sigma), 0.4 μg hFluc mRNA, 0.3 μl protease inhibitor (cOmplete, EDTA-free, Roche), 12 U RNAse inhibitor (Ribolock, Thermo Scientific), 6 μL S30 premix without amino acids (Promega), plus water to a final reaction volume of 15 μL were incubated for 1 h at 37°C. The reaction was stopped on ice before adding 75 μL of luciferase assay reagent (Promega) and recording of luminescence.

### 2.4. In-vitro antimicrobial susceptibility testing

Minimal inhibitory concentrations (MICs) of antibiotics were determined at four different study sites by broth microdilution assays according to the European Committee on Antimicrobial Susceptibility Testing (EUCAST) reference method ISO 20776-1:2019. In brief, antibiotics were serially twofold diluted in cation-adjusted Mueller-Hinton broth (CAMHB), inoculated to a final cell density of 10^6^ CFU/mL and incubated for 16-20 h at 35 ± 2°C followed by visual inspection for inhibition of bacterial growth. For the twenty CPE isolates, the susceptibility to gentamicin, tobramycin, and amikacin was determined according to the EUCAST disk diffusion method instead [19]. *E. coli* ATCC 25922 was used as quality control strain in all experimental setups.

### 2.5. Mutational apramycin resistance

Frequency of resistance experiments were performed by plating ∼10^9^ CFU on solid MH-II plates containing antibiotic concentrations corresponding to 4- or 8-times the agar-MIC for each strain. The plates were incubated at 37° C for 48 hours, and colonies were counted at 24 and 48 hours. Selected colonies were isolated on plates containing the same concentration of antibiotic. Colonies were prepared for whole genome sequencing using an Epicentre Masterpure DNA Preparation Kit (Nordic Biolabs, Sweden). Genomic DNA libraries were constructed using Nextera XT Library Preparation Kit (Illumina, San Diego, CA USA) and the libraries were sequenced on a MiSeq Device (Illumina, San Diego, CA USA). Sequence analysis was performed using CLC Genomics Workbench (Qiagen, Venlo, Netherlands) and the ResFinder Database [20-22].

### 2.6. Mouse peritonitis infection models

Mouse peritonitis-sepsis studies were performed as described previously [23]. In brief, procured NMRI female mice were acclimatized for a week before the study and allowed free access to chow and water. For dose-response studies, five to six weeks old mice weighing 26-30 g were acquired from Taconic (Denmark) and kept in cages of three. For all other studies, four weeks old mice weighing 18-22 g were acquired from Harlan-Stone (Netherlands) and kept in cages of eight. For each treatment and time point, three mice were used per group. Mice were inoculated intraperitoneally with 0.5 ×10^7^ CFU in 0.5 mL of sterile saline. One hour post inoculation, 0.2 ml of EBL-1003 in physiologic saline was injected subcutaneously. After mice were sacrificed, blood was obtained by cardiac puncture and collected into EDTA coated tubes. Peritoneal fluid was recovered after intraperitoneal injection of 2 ml of sterile saline and manual massage of the abdomen. Both blood and peritoneal fluid were immediately processed for quantitative counts by spread of undiluted or ten-fold dilutions on Mueller-Hinton or bromothymol (blue) agar plates. Mice in the dose response studies received dose levels between 0.39 and 200 mg/kg in twofold increments and were sacrificed five hours post inoculation. Mice in all other studies received a second dose four hours post inoculation, for a total dose of 2 × 80 mg/kg, and were sacrificed six hours post inoculation.

### 2.7. Statistics

For curvilinear analysis of antibiotic concentration versus effect (i.e. CFU counts *in vitro* or *in vivo*) the log-log sigmoid Hill-equation was applied using Graph-Pad Prism version 9.3.1 (GraphPad Software, Boston, MA, USA).

### 2.8. Ethics

All *in vivo* experiments were conducted under the supervision of the Danish Animal Ethical Council. Studies at SSI were conducted under license 2014-15-0201-00171. Studies at Rigshospitalet were conducted under license 2017–15-0201–01274.

## 3. Results

### 3.1. Aminoglycoside resistance gene annotations in the genomes of blood-culture isolates

First, we analyzed the resistance gene annotations of over 40 000 blood-culture isolates deposited in the National Database of Antibiotic Resistant Organisms (NDARO) to understand the prevalence and identity of aminoglycoside resistance genes. *Enterobacterales* represented the largest group with around 15 380 genomes and *K. pneumoniae* (ca. 50%) and *E. coli/Shigella* (ca. 38%) being the most abundant (**Fig. 1A-B**).

**Fig. 1.**
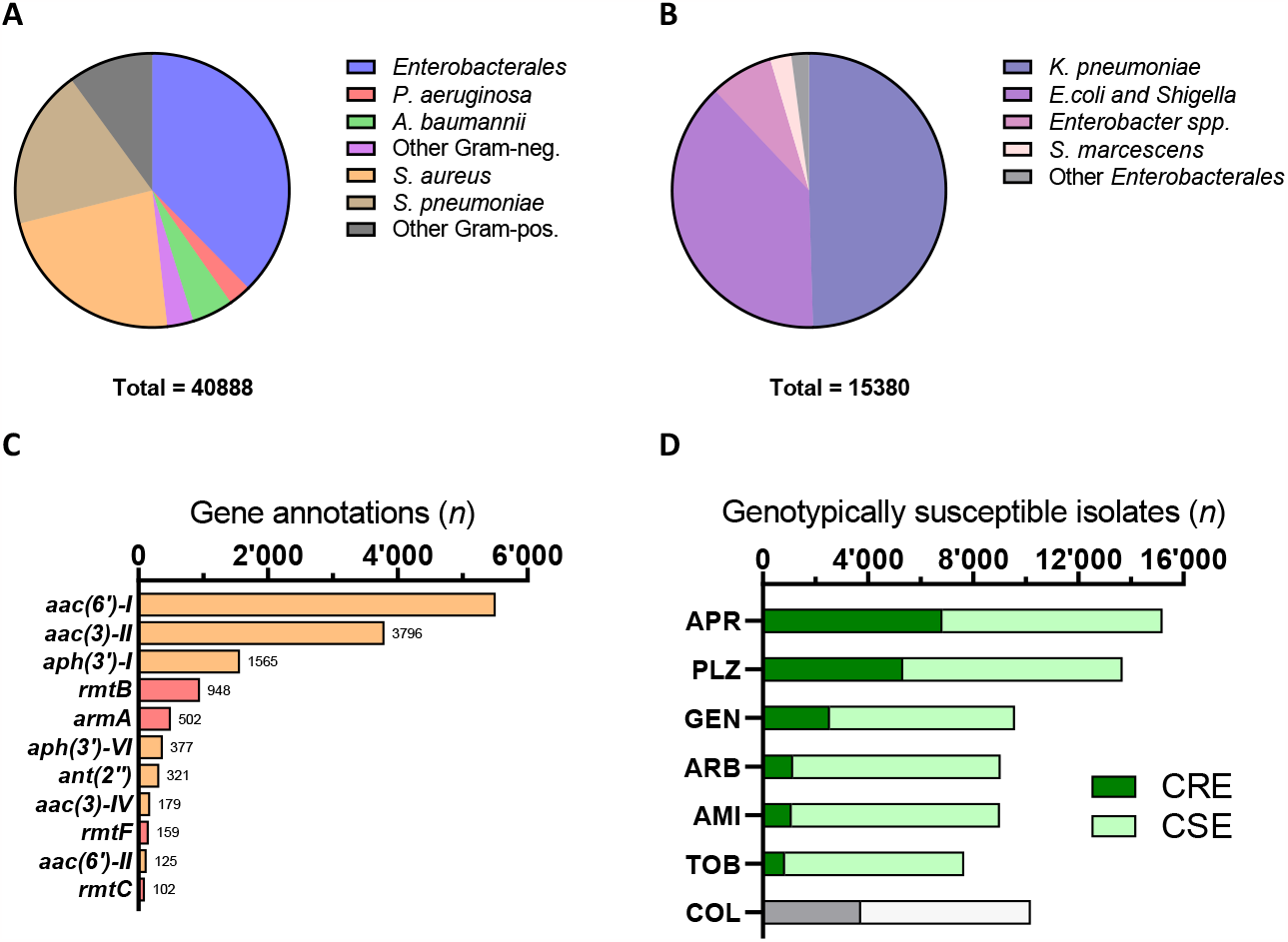
Data mining of clinical blood-culture isolate genomes deposited in the NDARO. (A) Distribution of bacterial pathogens among 40 888 blood-culture isolates deposited in the NDARO. (B) Species distribution within the 15 380 *Enterobacterales* blood culture isolates. (C) Prevalence of aminoglycoside resistance genes in the *Enterobacterales* blood-culture isolates. Aminoglycoside-modifying enzymes are shown in orange, 16S rRNA methylases are shown in red. (D) The genotypic susceptibility of the *Enterobacterales* blood culture isolates to apramycin (APR) in comparison to plazomicin (PLZ), amikacin (AMI), arbekacin (ARB), tobramycin (TOB), gentamicin (GEN), and colistin (COL) as determined by resistance gene annotations in 15 380 *Enterobacterales* genomes, including 6973 carbapenemase genotypes.

*aac(6’), aac(3)* and *aph(3’)* were found to be the most prevalent aminoglycoside resistance determinants in the blood-derived *Enterobacterales* isolates (**Fig. 1C**), followed by the 16S-rRNA methyltransferases (RMTases), which are often referred to as pan-aminoglycoside resistance genes due to conferring high-level resistance to all aminoglycoside antibiotics in clinical use including plazomicin. The aminoglycoside acetyltransferase gene *aac(3)-IV* was the only apramycin resistance gene present in the studied panel.

Knowledge of the phenotypic manifestation of each of the resistance genes (**Table S1**) can be used to predict differential drug susceptibilities suggesting 15 201 (99%) of drug-resistant *Enterobacterales* blood-culture isolates were susceptible to apramycin, compared to susceptibility rates < 90% for all other drugs studied (**Fig. 1D**). The difference between distinct aminoglycoside antibiotics is most pronounced in the carbapenemase subpopulation, with 98% of carbapenem-resistant *Enterobacterales* being genotypically susceptible to apramycin, as opposed to as low as 12-16% for tobramycin and amikacin (**Fig. 1D**).

### 3.2. Activity of aminoglycoside antibiotics against m^7^G1405-methylated ribosomes

Since RMTase genes appear to be a critical determinant for aminoglycoside resistance in blood-culture isolates, we decided to compare the inhibitory effect of various aminoglycoside antibiotics in a cell-free translation assay using m^7^G1405-methylated *E. coli* and *K. pneumonia* ribosomes. The IC_50_ of apramycin in inhibiting the synthesis of luciferase was determined as 0.03 and 0.04 μM, respectively. The activity of other deoxystreptamine-containing antibiotics was at least a thousand-fold lower, with IC_50_ values ranging from 42 μM (tobramycin) to > 800 μM (amikacin) in comparison (**Fig. 2A**).

**Fig. 2.**
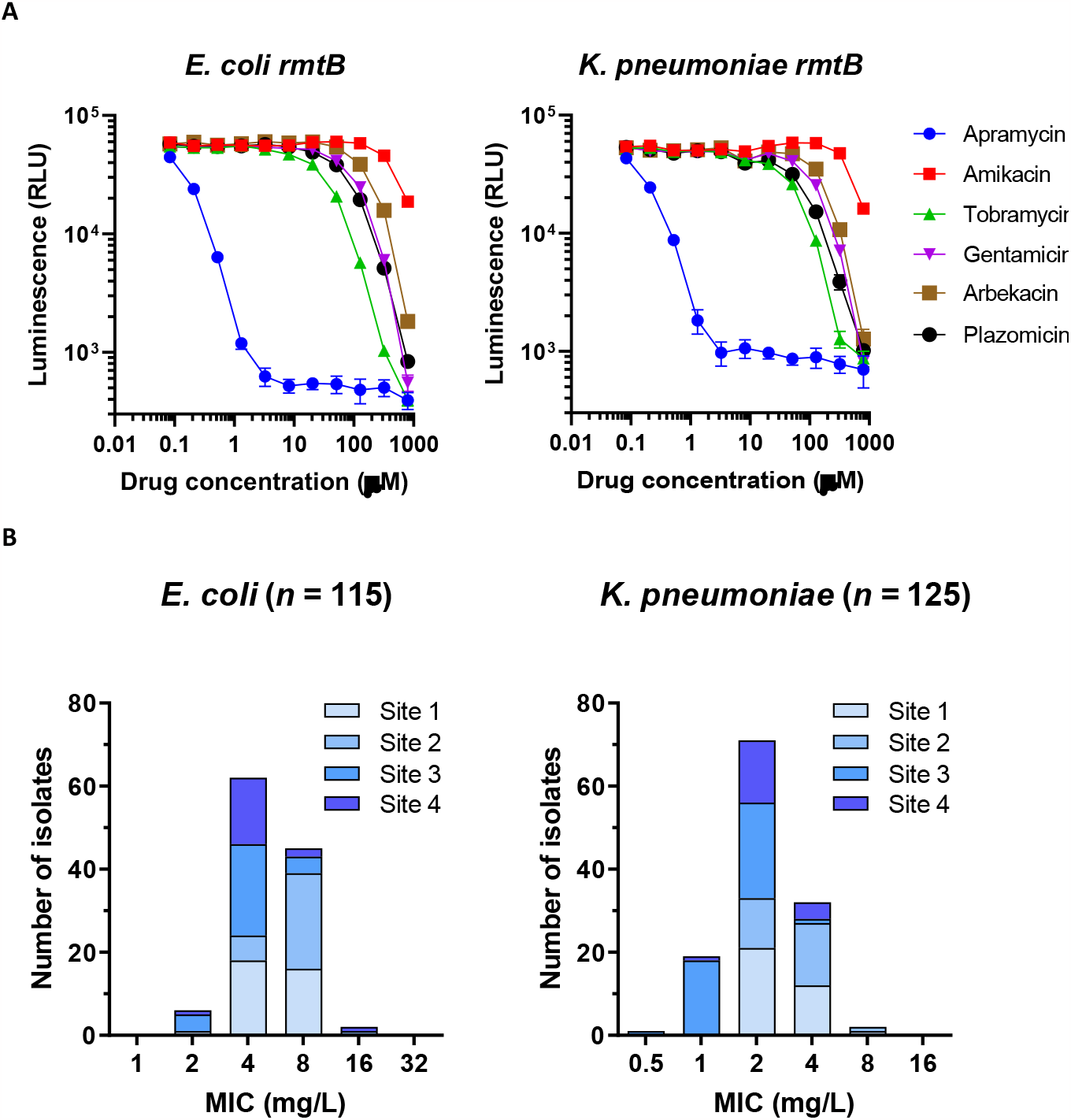
In-vitro activity of apramycin against *Escherichia coli* and *Klebsiella pneumoniae*. (A) In-vitro translation inhibition assay with S30 extracts of G1405-methylated *E. coli* or *K. pneumonia* isolates. (B) MIC distribution of susceptible wild-type bacterial isolates tested at 4 distinct centers (grey shadings) to determine a tentative epidemiological cutoff (ECOFF).

### 3.3. In vitro susceptibility of Escherichia coli and Klebsiella pneumoniae wild type isolates

Next, we performed a multicenter study on the activity of apramycin in broth microdilution assays with aminoglycoside-susceptible *E. coli* and *K. pneumoniae* clinical isolates. The MICs of apramycin against *E. coli* varied in a narrow Gaussian distribution from 2 to 16 mg/l (total of 4 dilution steps) with a modal MIC and MIC_50_ both at 4 mg/L, and an MIC_90_ of 8 mg/mL (**Fig. 2B**). The MIC distribution for *K. pneumoniae* was shifted to one dilution step lower in comparison, with a modal MIC and MIC_50_ both at 2 mg/L, an MIC_90_ of 4 mg/L, and within a range of 0.5 to 8 mg/L. Tentative ECOFFs were assumed to be 8 mg/mL for *E. coli* and 4 mg/L for *K. pneumoniae*, comprising 98.3 % and 98.4 % of isolates tested, respectively.

### 3.4. Frequency of mutational apramycin resistance

Aminoglycoside antibiotics bind to the 16S-rRNA of the bacterial ribosome where they inhibit protein synthesis. Since *Enterobacterales* carry multiple 16S rRNA genes, mutational resistance to aminoglycoside antibiotics is low and has not been described to be of clinical relevance. We determined the frequency of apramycin resistance for fifteen clinical isolates including aminoglycoside susceptible and aminoglycoside resistant phenotypes to find frequencies of ≤ 10^−8^ at 4 × MIC and ≤ 10^−10^ at 8 × MIC for most isolates (**Table S3**). Four of the resistant colonies were further characterized and found to be small colony variants with mutations in the *cydA* or *ubiD* respiratory pathway genes thought to be relevant for aminoglycoside uptake, but also causing large decreases in relative fitness (**Table S5**).

### 3.5. Aminoglycoside susceptibility of highly drug resistant blood-culture isolates

Next, we tested the activity of apramycin in comparison to other aminoglycosides against highly drug resistant clinical isolates comprising several metallo-β-lactamases, KPC’s, various OXA-, SHV-, CTX-M- and CMY-genes (**Table S4**). In addition to these carbapenemase genes, all isolates carried genes for several other β-lactamases rendering them resistant towards a broad range of penicillins and cephalosporins. Various aminoglycoside-resistance genotypes were also present in these isolates. Only one strain, KP3, carried an *aac(3)-IVa*-like gene that conferred apramycin resistance (**Table S4**). All other isolates were susceptible to apramycin in spite of the presence of multiple and diverse aminoglycoside resistance genes, with MICs of 4-8 mg/L for *E. coli* and 2-4 mg/L for *K. pneumoniae* equivalent to the wild-type MIC distribution reported above. Amikacin was the most active of the clinical care aminoglycosides with 12 of 20 isolates found to be amikacin resistant, as opposed to 16 of 20 isolates found resistant to both gentamicin and tobramycin (**Table S4**).

### 3.6. Apramycin efficacy and dose response in a mouse sepsis model

Proof of concept for the efficacy of apramycin (EBL-1003) in the treatment of blood stream infections was established with a mouse sepsis-peritonitis model using *E. coli* ATCC 25922 (MIC, 4 mg/L) and *K. pneumoniae* SSI 3010 (MIC, 2 mg/L) reference strains. The maximal killing effect (E_max_) of EBL-1003 for the two pathogens corresponded to 4.2 - 4.5 log CFU reduction in blood and 3.5 - 3.7 log in peritoneum (**Fig. 3**). The *E. coli* infection required a dose of 7 - 9 mg/kg for stasis and 12 - 13 mg/kg for a 1-log CFU reduction in both compartments. The corresponding doses were 1 - 3 mg/kg and 5 - 9 mg/kg, respectively, for the *K. pneumoniae* infection, reflecting the differences in MIC for the two species (**Table 1**). However, since the slope of the dose-response curves in the two compartments was steeper for *E. coli* than for *K. pneumoniae*, the ED90 dose of 18-24 mg/kg was lower than those for *K. pneumoniae* at > 50 mg/kg.

**Table 1.** Dose-response regression analysis.

|  | *E. coli* ATCC 25922 |  | *K. pneumoniae* SSI 3010 |  |
| --- | --- | --- | --- | --- |
|  | Blood | Peritoneum | Blood | Peritoneum |
| Stasis | 8.4 | 7.4 | 2.6 | 1.3 |
| -1 log | 13.1 | 11.8 | 9.4 | 4.8 |
| -2 log | 29.0 | NA<sup>1</sup> | 29.5 | 15.6 |
| ED<sub>50</sub> | 9.2 | 7.8 | 12.7 | 5.4 |
| ED<sub>90</sub> | 24.1 | 17.8 | > 200 | 52.0 |
| R<sup>2</sup> | 0.85 | 0.96 | 0.94 | 0.91 |

<sup>1</sup> NA, not applicable, CFU count reduction < 2 log

**Fig. 3.**
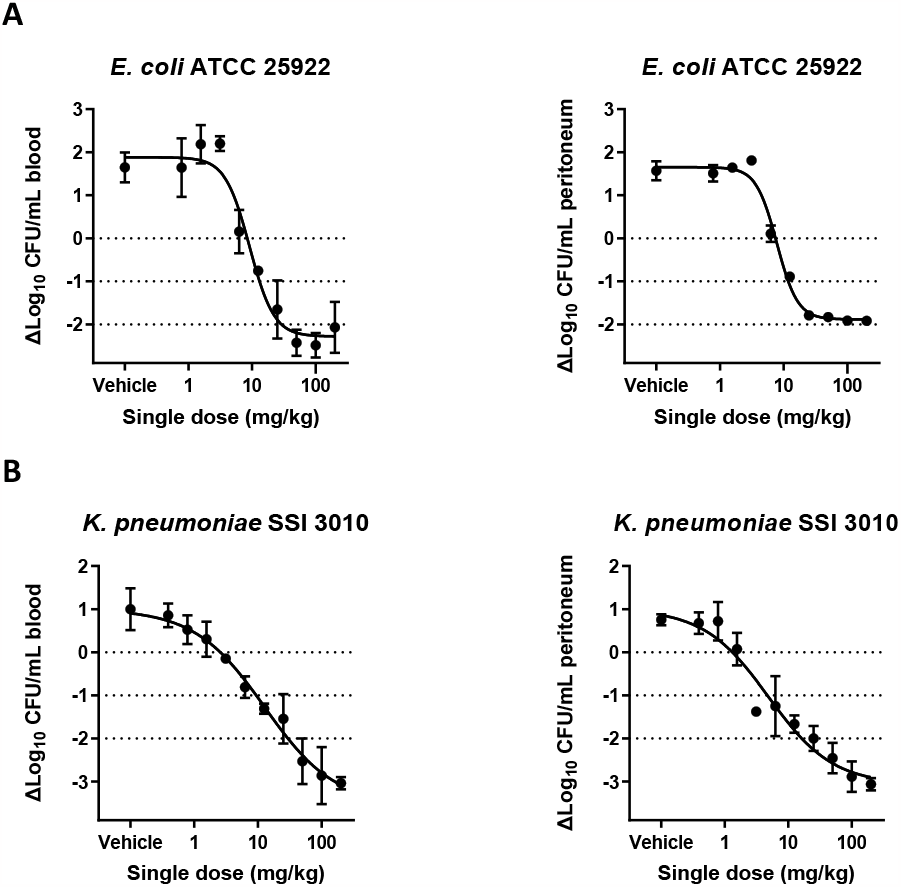
Efficacy of EBL-1003 in a murine dose-range peritonitis model. (**A**) Dose-range response study with *E. coli* ATCC 25922, MIC = 4 μg/mL, *n* = 3 animals per dose group. (**B**) Dose-range response study with *K. pneumonia* strain SSI 3010, MIC = 1-2 μg/mL, *n* = 3 animals per dose group. Results of the regression analysis are summarized in **Table 1**.

### 3.7. Apramycin efficacy against highly drug resistant blood-culture isolates

Lastly, we wanted to test whether the potent *in vitro* activity of apramycin against highly drug resistant *Enterobacterales* isolates reported above would translate into equally potent *in vivo* efficacy against blood stream infections with carbapenem-resistant *Enterobacterales*, a group of pathogens that has proven particularly difficult to treat and therefore represents a very high medical need. Pathogens for infection were selected from the twenty isolates described above to represent a diversity of carbapenemase genotypes and based on their confirmed resistance to carbapenems and all aminoglycoside antibiotics except apramycin. Based on the dose-response results above, a dose of 2 × 80 mg/kg was selected so as to result in at least 3-log CFU reduction in both peritoneum and blood, and add up to a total dose that would result in an AUC similar to the AUC observed in humans after a single dose of 30 mg/kg [24].

EBL-1003 fully resolved bacteremia of > 10^5^ CFU/mL within ≤ 5 hours to below the CFU detection limit for all of the three *E. coli* and five *K. pneumoniae* isolates studied, including all carbapenem- and pan-aminoglycoside resistant isolates (**Fig. 4**). Bacterial burden in the peritoneum was fully resolved for four of the five *K. pneumoniae* isolates, and reduced by at least three logs for the other four isolates. Administration of 2 × 80 mg/kg of amikacin did not reduce the CFU counts when compared to the vehicle control mice, as it was expected based on their amikacin-resistant phenotype confirmed *in vitro* (**Table S4**).

**Fig. 4.**
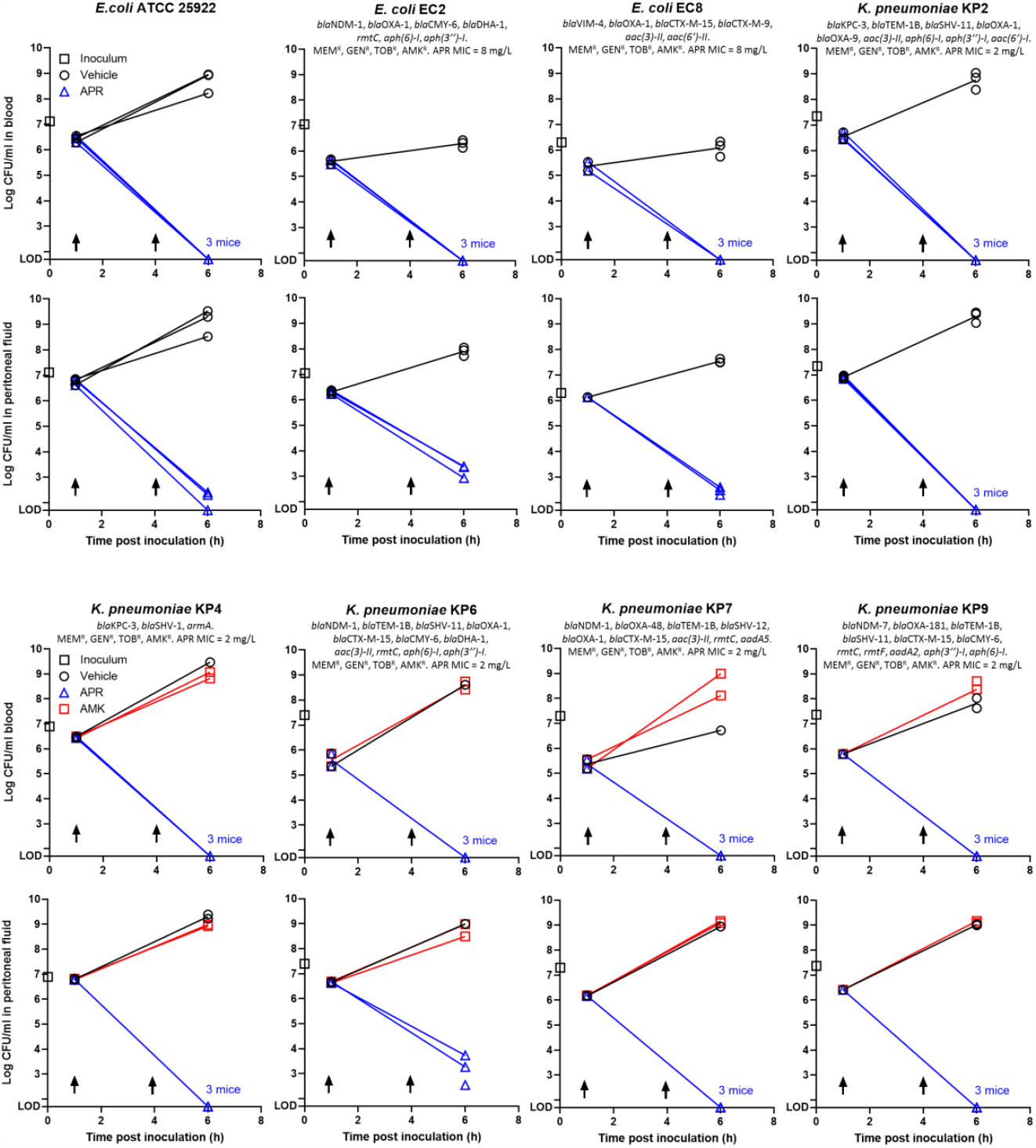
Efficacy of EBL-1003 in the murine sepsis-peritonitis model. The figures show CFU counts in blood and peritoneum for each isolate tested. Data points for t_0_ indicate the CFU count of the inoculum. Arrows indicate subcutaneous treatment with 80 mg/kg apramycin. Each data point denotes result for one mouse; where points coalesce with same result, text indicates how many mice were involved. Resistance genes for β-lactamases and aminoglycoside resistance are shown for each strain, as well as the MIC for apramycin.

## 4. Discussion

The results of this study suggest apramycin provides excellent coverage and efficacy in the treatment of bloodstream infections caused by multi-drug resistant *Enterobacterales*. These findings are in line with previous reports of EBL-1003, a pharmaceutical grade preparation of apramycin, demonstrating encouraging efficacy against Gram-negative bacilli in thigh, lung, and urinary tract infection models, and against *Staphylococcus aureus* in a septicemia model [10-14]. The in-vitro coverage of several hundred *Enterobacterales* and blood-culture isolates has been demonstrated recently [25].

EBL-1003 has been developed due to the rapidly increasing resistance world-wide, especially in the most important pathogens *E. coli* and *K. pneumoniae*, which rank high on all lists of clinical burden or cost of resistance [26-28]. The genotypic susceptibility to apramycin when screening the NDARO database with most bacteremia pathogens represented by *E. coli* and *K. pneumoniae* revealed an overwhelming coverage by apramycin in comparison to other aminoglycosides in clinical use including plazomicin. This in-silico finding is in agreement with recent reports on the in-vitro data for several hundred blood-culture isolates [25, 29], except that it derives its power from an unprecedented larger number of genomes from around the world [16]. The only apramycin resistance gene identified in the 15 380 *Enterobacterales* blood-culture isolates was *aac(3)-IV*, which is in support of earlier reports of this gene being the only resistance gene of clinical relevance to apramycin [30]. However, it’s incidence in the studied population (1.1%) was at least tenfold lower than that of other aminoglycoside resistance genes such as RMTases (11.1%), *aac(3)-II* (24.5%), or *aac(6’)-I* (35.8%), awarding best-in-class genotypic coverage. The breadth of such genotypic analyses also provides valuable context to another study that reported higher incidence of *aac(3)-IV* when studying a selected population of *K. pneumoniae* ST258 strains [31].

Apramycin MICs for wild type *E. coli* and *K. pneumoniae* lie within a narrow range from 0.5 to 16 mg/L, with modal MICs of 4 mg/L for *E. coli* and 2 mg/L for *K. pneumoniae* (**Fig. 2B**). The slightly better activity against *Klebsiella*, i.e. lower MICs, than for *E. coli* is seen also for amikacin and tobramycin with MICs around 1-2 mg/L for both aminoglycosides, while gentamicin shows a median MIC of 0.5 mg/L for both pathogens (https://mic.eucast.org). ECOFFs have been tentatively determined as 8 mg/L and 4 mg/L for *E. coli* and *K. pneumoniae*, respectively. EBL-1003 showed remarkably good activity against the collection of MDR CPEs which in addition to carbapenemase genes harbored several aminoglycoside resistance genes (1-5 in each strain); while mostly resistant towards the standard aminoglycosides, all strains except one, KP3 with an *aac(3)-IV* gene, were susceptible to EBL-1003. This activity was confirmed in the in vivo efficacy study (see **Fig. 3** and **Fig. 4**) where the drug eradicated the bacteria from blood, and in most cases from the peritoneum with Log 1-2 bacteria remaining after 5 h in the peritoneum for KP6, EC2 and EC8 (**Fig. S1**). Thus, EBL-1003 was highly effective in clearing the blood stream in the mouse sepsis/peritonitis model, but the activity in the peritoneal fluid appeared to be dependent on the MIC of the infecting pathogen. *E. coli* isolates with MICs of 8 mg/L for EBL-1003 were apparently more difficult to eradicate from the peritoneum than *K. pneumoniae* with MICs of 2 mg/L. The reason for the difference in response curves between *E. coli* and *K. pneumoniae* in the dose ranging study (**Fig. 3**) is not clear, but *K. pneumoniae* are more susceptible, i.e. lower MICs, than for *E. coli* and this is reflected in the lower doses needed for effect against *Klebsiella* than for *E. coli*. The doses chosen for the efficacy study were based on the dose-ranging study in this paper but also on previous studies, which have tried to calculate the dose needed for clinical use in humans [13]. The optimal dose suggested for humans has been proposed as 30 mg/kg resulting in an AUC of about 700 μmol*h/L [13, 15, 24]. The 2 × 80 mg/kg dose used in this study results in a similar AUC in mice and a C_max_ of about 150-200 mg/L, which would ensure 19-25 times the ECOFF for *E. coli*, i.e. a promising PKPD index for treatment with this drug class [32].

The limitations of this study may lie in the relatively low number of strains tested and evaluated for a tentative ECOFF and in the animal studies. On the other hand, the variation in the MICs among the wild type organisms was so low, that there is no reason to believe that adding more strains would change the results. The genotypic susceptibility screen based on the NDARO database, albeit a useful surrogate powered by a very large sample size, has its limitations, too. It has previously been shown, that there may be a mismatch between genotype and phenotype, notably that presence of a resistance gene does not correspond to phenotypic resistance [33, 34]. On the other hand, all aminoglycoside resistance genes found in the CPE isolates corresponded 100% with resistance or susceptibility of apramycin and standard aminoglycosides.

In conclusion, this study confirms that apramycin is a very interesting and important antibiotic for further clinical development: 1. apramycin is very effective *in vitro* with MICs against wild-type *E. coli* and *K. pneumoniae* very close to the MICs of standard aminoglycosides. 2. apramycin is active against a large range of recent *Enterobacterales* isolates from all over the world harboring a wide range of aminoglycoside resistance genes; the only gene conferring resistance towards apramycin i.e. *aac(3)-IV* is very infrequent or rare among clinical isolates. 3. apramycin has a very low potential for developing mutational resistance, which is also the case for standard aminoglycosides. 4. apramycin formulation EBL-1003 was very effective in clearing the blood stream and the peritoneum during experimental infection with MDR carbapenemase-producing *K. pneumoniae* which were resistant towards amikacin, and amikacin on the same doses had no effect in this model.

Taken together with the lower oto- and nephrotoxic potential of EBL-1003, this compound therefore has huge clinical potential and support for its further development should be obvious.

## Ethical approval

All in vivo experiments were conducted under the supervision of the Danish Animal Ethical Council. Studies were conducted under licenses 2014-15-0201-00171 and 2017–15-0201–01274.

## Supplementary materials

Supplementary naterial associated with this article is available in the online version.

## Acknowledgements

The authors are grateful to Frank Hansen at Statens Serum Institute for providing carbapenem-resistant clinical isolates for this study.

