## Supplementary Tables and Figures for "Apramycin efficacy in the treatment of carbapenem-resistant *Enterobacterales* in murine blood stream infection models"

**Table S1.** Resistance genes of the NDARO Gene Catalog used in the data base analysis

| Drug | Associated resistance genes |
|---|---|
| Apramycin | *aac(3)-IV, apmA, npmA, kamB* |
| Amikacin | *aac(6')-I, aph(3'), ant(4'), aadD, npmA, armA, rmtA, rmtB, rmtC, rmtD, rmtE, rmtF, rmtG, rmtH* |
| Tobramycin | *aac(6')-I, aac(6')-II, aac(3)-II, aac(3)-III, aac(3)-IV, aac(2'), aph(2''), ant(2''), ant(4'), aacA35, aadD, npmA, armA, rmtA, rmtB, rmtC, rmtD, rmtE, rmtF, rmtG, rmtH* |
| Gentamicin | *aac(6')-II, aac(6')-Ib', aac(6')-Ib4, aac(6')-Ib11, aacA35, aac(3)-I, aph(2''), ant(2''), npmA, armA, rmtA, rmtB, rmtC, rmtD, rmtE, rmtF, rmtG, rmtH* |
| Arbekacin | *aac(6')-I, aacA35, npmA, armA, rmtA, rmtB, rmtC, rmtD, rmtE, rmtF, rmtG, rmtH* |
| Plazomicin | *aac(2'), aph(2''), npmA, armA, rmtA, rmtB, rmtC, rmtD, rmtE, rmtF, rmtG, rmtH* |
| Colistin | *almE, almF, almG, arnC, basR, colR, colS, cprS, crrB, icr-Mo, lapB, lpxA, lpxC, lpxM, mcr-1, mcr-2, mcr-3, mcr-4, mcr-5, mcr-6, mcr-7, mcr-8, mcr-9, mcr-10, nfxB, parS, phoP, phoQ, pmrA, pmrB, pmrC* |
| Carbapenems | *amgS, amrR, bla2, blaADC, blaAFM, blaAIM, blaALG11, blaALG6, blaALI, blaANA, blaAXC, blab, blaB1PEDO, blaB3PEDO, blaBIC, blaBJP, blaBKC, blaCAM, blaCAR, blaCAU, blaCPS, blaCRD3, blaCRH, blaCRP, blaCVI, blaDHT2, blaDIM, blaEAM, blaEBR, blaECM, blaECV, blaEFM, blaELM, blaESP, blaEVM, blaFEZ, blaFIA, blaFIM, blaFPH, blaFRI, blaGES, blaGIM, blaGMB, blaGOB, blaGPC, blaGRD23, blaGRD33, blaHMB, blaIMI, blaIMP, blaIND, blaJOHN, blaKHM, blaKPC, blaL1, blaLMB, blaMAB, blaMOC, blaMSI, blaMUS, blaMYO, blaMYX, blaNDM, blaNWM, blaORR, blaOXA, blaPAD, blaPAM, blaPFM, blaPJM, blaPLN, blaPNGM, blaPOM, blaPST, blaRAD, blaRm3, blaRSA2, blaSFC, blaSFH, blaSHD, blaSHN, blaSHW, blaSIM, blaSMB, blaSME, blaSPG, blaSPM, blaSPN79, blaSPS, blaSTA, blaTHIN-B, blaTMB, blaTTU, blaTUS, blaVAM, blaVCC, blaVIM, blaVMB, blaWUS, blaYEM, blaZOG, bpeR, cfiA, cphA, cphA1, crxA, czcS, ftsI, mexR, nalC, ompK35, ompK36, oprD, parS, varG.* |

**Table S2. Bacterial isolates used to determine a tentative epidemiologic cutoff (ECOFF).**

| Species | Phenoytpe | Isolate | Site | ID | Isolate ID | APR MIC (µg/mL) |
|---|---|---|---|---|---|---|
| *Escherichia coli* | Aminoglycoside susceptible | 1 | Site 1 | UZH | 1 | 8 |
| *Escherichia coli* | Aminoglycoside susceptible | 2 | Site 1 | UZH | 2 | 4 |
| *Escherichia coli* | Aminoglycoside susceptible | 3 | Site 1 | UZH | 3 | 8 |
| *Escherichia coli* | Aminoglycoside susceptible | 4 | Site 1 | UZH | 4 | 4 |
| *Escherichia coli* | Aminoglycoside susceptible | 5 | Site 1 | UZH | 5 | 8 |
| *Escherichia coli* | Aminoglycoside susceptible | 6 | Site 1 | UZH | 6 | 8 |
| *Escherichia coli* | Aminoglycoside susceptible | 7 | Site 1 | UZH | 7 | 4 |
| *Escherichia coli* | Aminoglycoside susceptible | 8 | Site 1 | UZH | 8 | 4 |
| *Escherichia coli* | Aminoglycoside susceptible | 9 | Site 1 | UZH | 9 | 4 |
| *Escherichia coli* | Aminoglycoside susceptible | 10 | Site 1 | UZH | 10 | 8 |
| *Escherichia coli* | Aminoglycoside susceptible | 11 | Site 1 | UZH | 11 | 8 |
| *Escherichia coli* | Aminoglycoside susceptible | 12 | Site 1 | UZH | 12 | 4 |
| *Escherichia coli* | Aminoglycoside susceptible | 13 | Site 1 | UZH | 13 | 8 |
| *Escherichia coli* | Aminoglycoside susceptible | 14 | Site 1 | UZH | 14 | 4 |
| *Escherichia coli* | Aminoglycoside susceptible | 15 | Site 1 | UZH | 15 | 4 |
| *Escherichia coli* | Aminoglycoside susceptible | 16 | Site 1 | UZH | 16 | 8 |
| *Escherichia coli* | Aminoglycoside susceptible | 17 | Site 1 | UZH | 17 | 4 |
| *Escherichia coli* | Aminoglycoside susceptible | 18 | Site 1 | UZH | 18 | 8 |
| *Escherichia coli* | Aminoglycoside susceptible | 19 | Site 1 | UZH | 19 | 4 |
| *Escherichia coli* | Aminoglycoside susceptible | 20 | Site 1 | UZH | 20 | 4 |
| *Escherichia coli* | Aminoglycoside susceptible | 21 | Site 1 | UZH | 21 | 8 |
| *Escherichia coli* | Aminoglycoside susceptible | 22 | Site 1 | UZH | 22 | 4 |
| *Escherichia coli* | Aminoglycoside susceptible | 23 | Site 1 | UZH | 23 | 8 |
| *Escherichia coli* | Aminoglycoside susceptible | 24 | Site 1 | UZH | 24 | 4 |
| *Escherichia coli* | Aminoglycoside susceptible | 25 | Site 1 | UZH | 25 | 4 |
| *Escherichia coli* | Aminoglycoside susceptible | 26 | Site 1 | UZH | 26 | 4 |
| *Escherichia coli* | Aminoglycoside susceptible | 27 | Site 1 | UZH | 27 | 8 |
| *Escherichia coli* | Aminoglycoside susceptible | 28 | Site 1 | UZH | 28 | 4 |
| *Escherichia coli* | Aminoglycoside susceptible | 29 | Site 1 | UZH | 29 | 8 |
| *Escherichia coli* | Aminoglycoside susceptible | 30 | Site 1 | UZH | 30 | 2 |
| *Escherichia coli* | Aminoglycoside susceptible | 31 | Site 1 | UZH | 31 | 8 |
| *Escherichia coli* | Aminoglycoside susceptible | 32 | Site 1 | UZH | 32 | 4 |
| *Escherichia coli* | Aminoglycoside susceptible | 33 | Site 1 | UZH | 33 | 8 |
| *Escherichia coli* | Aminoglycoside susceptible | 34 | Site 1 | UZH | 34 | 4 |
| *Escherichia coli* | Aminoglycoside susceptible | 35 | Site 1 | UZH | 35 | 8 |
| *Escherichia coli* | Aminoglycoside susceptible | 36 | Site 2 | RH | 1 | 8 |
| *Escherichia coli* | Aminoglycoside susceptible | 37 | Site 2 | RH | 2 | 8 |
| *Escherichia coli* | Aminoglycoside susceptible | 38 | Site 2 | RH | 3 | 4 |
| *Escherichia coli* | Aminoglycoside susceptible | 39 | Site 2 | RH | 4 | 8 |
| *Escherichia coli* | Aminoglycoside susceptible | 40 | Site 2 | RH | 5 | 4 |
| *Escherichia coli* | Aminoglycoside susceptible | 41 | Site 2 | RH | 6 | 8 |
| *Escherichia coli* | Aminoglycoside susceptible | 42 | Site 2 | RH | 7 | 4 |
| *Escherichia coli* | Aminoglycoside susceptible | 43 | Site 2 | RH | 8 | 8 |
| *Escherichia coli* | Aminoglycoside susceptible | 44 | Site 2 | RH | 125 | 4 |
| *Escherichia coli* | Aminoglycoside susceptible | 45 | Site 2 | RH | 126 | 4 |
| *Escherichia coli* | Aminoglycoside susceptible | 46 | Site 2 | RH | 128 | 8 |
| *Escherichia coli* | Aminoglycoside susceptible | 47 | Site 2 | RH | 129 | 8 |
| *Escherichia coli* | Aminoglycoside susceptible | 48 | Site 2 | RH | 130 | 8 |
| *Escherichia coli* | Aminoglycoside susceptible | 49 | Site 2 | RH | 131 | 8 |
| *Escherichia coli* | Aminoglycoside susceptible | 50 | Site 2 | RH | 132 | 16 |
| *Escherichia coli* | Aminoglycoside susceptible | 51 | Site 2 | RH | FE01 | 8 |
| *Escherichia coli* | Aminoglycoside susceptible | 52 | Site 2 | RH | FE02 | 8 |
| *Escherichia coli* | Aminoglycoside susceptible | 53 | Site 2 | RH | FE03 | 4 |
| *Escherichia coli* | Aminoglycoside susceptible | 54 | Site 2 | RH | FE04 | 8 |
| *Escherichia coli* | Aminoglycoside susceptible | 55 | Site 2 | RH | 2LT | 8 |
| *Escherichia coli* | Aminoglycoside susceptible | 56 | Site 2 | RH | 4LT | 8 |
| *Escherichia coli* | Aminoglycoside susceptible | 57 | Site 2 | RH | 10LT | 8 |
| *Escherichia coli* | Aminoglycoside susceptible | 58 | Site 2 | RH | 13LT | 8 |
| *Escherichia coli* | Aminoglycoside susceptible | 59 | Site 2 | RH | 22LT | 8 |
| *Escherichia coli* | Aminoglycoside susceptible | 60 | Site 2 | RH | 23LT | 8 |
| *Escherichia coli* | Aminoglycoside susceptible | 61 | Site 2 | RH | 30LT | 8 |
| *Escherichia coli* | Aminoglycoside susceptible | 62 | Site 2 | RH | 35LT | 8 |
| *Escherichia coli* | Aminoglycoside susceptible | 63 | Site 2 | RH | 36LT | 8 |

**Table S2** (continued).

| Species | Phenoytpe | Isolate | Site | ID | Isolate ID | APR MIC (μg/mL) |
|---|---|---|---|---|---|---|
| *Escherichia coli* | Aminoglycoside susceptible | 64 | Site 2 | RH | 37LT | 8 |
| *Escherichia coli* | Aminoglycoside susceptible | 65 | Site 2 | RH | 41LT | 8 |
| *Escherichia coli* | Aminoglycoside susceptible | 66 | Site 3 | UU | EN0030 | 4 |
| *Escherichia coli* | Aminoglycoside susceptible | 67 | Site 3 | UU | EN0031 | 4 |
| *Escherichia coli* | Aminoglycoside susceptible | 68 | Site 3 | UU | EN0034 | 8 |
| *Escherichia coli* | Aminoglycoside susceptible | 69 | Site 3 | UU | EN0035 | 4 |
| *Escherichia coli* | Aminoglycoside susceptible | 70 | Site 3 | UU | EN0036 | 4 |
| *Escherichia coli* | Aminoglycoside susceptible | 71 | Site 3 | UU | EN0038 | 4 |
| *Escherichia coli* | Aminoglycoside susceptible | 72 | Site 3 | UU | EN0040 | 4 |
| *Escherichia coli* | Aminoglycoside susceptible | 73 | Site 3 | UU | EN0041 | 4 |
| *Escherichia coli* | Aminoglycoside susceptible | 74 | Site 3 | UU | EN0042 | 4 |
| *Escherichia coli* | Aminoglycoside susceptible | 75 | Site 3 | UU | EN0044 | 2 |
| *Escherichia coli* | Aminoglycoside susceptible | 76 | Site 3 | UU | EN0045 | 4 |
| *Escherichia coli* | Aminoglycoside susceptible | 77 | Site 3 | UU | EN0046 | 4 |
| *Escherichia coli* | Aminoglycoside susceptible | 78 | Site 3 | UU | EN0047 | 4 |
| *Escherichia coli* | Aminoglycoside susceptible | 79 | Site 3 | UU | EN0048 | 4 |
| *Escherichia coli* | Aminoglycoside susceptible | 80 | Site 3 | UU | EN0049 | 4 |
| *Escherichia coli* | Aminoglycoside susceptible | 81 | Site 3 | UU | EN0050 | 4 |
| *Escherichia coli* | Aminoglycoside susceptible | 82 | Site 3 | UU | EN0051 | 2 |
| *Escherichia coli* | Aminoglycoside susceptible | 83 | Site 3 | UU | EN0052 | 4 |
| *Escherichia coli* | Aminoglycoside susceptible | 84 | Site 3 | UU | EN0053 | 2 |
| *Escherichia coli* | Aminoglycoside susceptible | 85 | Site 3 | UU | EN0054 | 4 |
| *Escherichia coli* | Aminoglycoside susceptible | 86 | Site 3 | UU | EN0055 | 8 |
| *Escherichia coli* | Aminoglycoside susceptible | 87 | Site 3 | UU | EN0056 | 4 |
| *Escherichia coli* | Aminoglycoside susceptible | 88 | Site 3 | UU | EN0057 | 8 |
| *Escherichia coli* | Aminoglycoside susceptible | 89 | Site 3 | UU | EN0058 | 8 |
| *Escherichia coli* | Aminoglycoside susceptible | 90 | Site 3 | UU | EN0059 | 2 |
| *Escherichia coli* | Aminoglycoside susceptible | 91 | Site 3 | UU | EN0060 | 4 |
| *Escherichia coli* | Aminoglycoside susceptible | 92 | Site 3 | UU | EN0061 | 4 |
| *Escherichia coli* | Aminoglycoside susceptible | 93 | Site 3 | UU | EN0062 | 4 |
| *Escherichia coli* | Aminoglycoside susceptible | 94 | Site 3 | UU | EN0154 | 4 |
| *Escherichia coli* | Aminoglycoside susceptible | 95 | Site 3 | UU | EN0155 | 4 |
| *Escherichia coli* | Aminoglycoside susceptible | 96 | Site 4 | IHMA | 1987026 | 4 |
| *Escherichia coli* | Aminoglycoside susceptible | 97 | Site 4 | IHMA | 1995158 | 4 |
| *Escherichia coli* | Aminoglycoside susceptible | 98 | Site 4 | IHMA | 2052714 | 4 |
| *Escherichia coli* | Aminoglycoside susceptible | 99 | Site 4 | IHMA | 2031478 | 4 |
| *Escherichia coli* | Aminoglycoside susceptible | 100 | Site 4 | IHMA | 2039005 | 4 |
| *Escherichia coli* | Aminoglycoside susceptible | 101 | Site 4 | IHMA | 1971705 | 8 |
| *Escherichia coli* | Aminoglycoside susceptible | 102 | Site 4 | IHMA | 1984591 | 4 |
| *Escherichia coli* | Aminoglycoside susceptible | 103 | Site 4 | IHMA | 1957336 | 4 |
| *Escherichia coli* | Aminoglycoside susceptible | 104 | Site 4 | IHMA | 2019232 | 4 |
| *Escherichia coli* | Aminoglycoside susceptible | 105 | Site 4 | IHMA | 2013839 | 8 |
| *Escherichia coli* | Aminoglycoside susceptible | 106 | Site 4 | IHMA | 1970605 | 4 |
| *Escherichia coli* | Aminoglycoside susceptible | 107 | Site 4 | IHMA | 1993665 | 4 |
| *Escherichia coli* | Aminoglycoside susceptible | 108 | Site 4 | IHMA | 1986506 | 4 |
| *Escherichia coli* | Aminoglycoside susceptible | 109 | Site 4 | IHMA | 2331987 | 2 |
| *Escherichia coli* | Aminoglycoside susceptible | 110 | Site 4 | IHMA | 2029891 | 4 |
| *Escherichia coli* | Aminoglycoside susceptible | 111 | Site 4 | IHMA | 2213479 | 4 |
| *Escherichia coli* | Aminoglycoside susceptible | 112 | Site 4 | IHMA | 2008120 | 4 |
| *Escherichia coli* | Aminoglycoside susceptible | 113 | Site 4 | IHMA | 1983050 | 16 |
| *Escherichia coli* | Aminoglycoside susceptible | 114 | Site 4 | IHMA | 1994117 | 4 |
| *Escherichia coli* | Aminoglycoside susceptible | 115 | Site 4 | IHMA | 2005104 | 4 |

**Table S2** (continued).

| Species | Phenoytpe | Isolate | Site | ID | Isolate ID | APR MIC (µg/mL) |
|---|---|---|---|---|---|---|
| *Klebsiella pneumoniae* | Aminoglycoside susceptible | 1 | Site 1 | UZH | 101 | 4 |
| *Klebsiella pneumoniae* | Aminoglycoside susceptible | 2 | Site 1 | UZH | 102 | 2 |
| *Klebsiella pneumoniae* | Aminoglycoside susceptible | 3 | Site 1 | UZH | 103 | 2 |
| *Klebsiella pneumoniae* | Aminoglycoside susceptible | 4 | Site 1 | UZH | 104 | 4 |
| *Klebsiella pneumoniae* | Aminoglycoside susceptible | 5 | Site 1 | UZH | 105 | 2 |
| *Klebsiella pneumoniae* | Aminoglycoside susceptible | 6 | Site 1 | UZH | 106 | 4 |
| *Klebsiella pneumoniae* | Aminoglycoside susceptible | 7 | Site 1 | UZH | 107 | 4 |
| *Klebsiella pneumoniae* | Aminoglycoside susceptible | 8 | Site 1 | UZH | 108 | 2 |
| *Klebsiella pneumoniae* | Aminoglycoside susceptible | 9 | Site 1 | UZH | 109 | 2 |
| *Klebsiella pneumoniae* | Aminoglycoside susceptible | 10 | Site 1 | UZH | 110 | 2 |
| *Klebsiella pneumoniae* | Aminoglycoside susceptible | 11 | Site 1 | UZH | 111 | 2 |
| *Klebsiella pneumoniae* | Aminoglycoside susceptible | 12 | Site 1 | UZH | 112 | 2 |
| *Klebsiella pneumoniae* | Aminoglycoside susceptible | 13 | Site 1 | UZH | 113 | 2 |
| *Klebsiella pneumoniae* | Aminoglycoside susceptible | 14 | Site 1 | UZH | 114 | 2 |
| *Klebsiella pneumoniae* | Aminoglycoside susceptible | 15 | Site 1 | UZH | 115 | 4 |
| *Klebsiella pneumoniae* | Aminoglycoside susceptible | 16 | Site 1 | UZH | 116 | 2 |
| *Klebsiella pneumoniae* | Aminoglycoside susceptible | 17 | Site 1 | UZH | 117 | 2 |
| *Klebsiella pneumoniae* | Aminoglycoside susceptible | 18 | Site 1 | UZH | 118 | 2 |
| *Klebsiella pneumoniae* | Aminoglycoside susceptible | 19 | Site 1 | UZH | 119 | 4 |
| *Klebsiella pneumoniae* | Aminoglycoside susceptible | 20 | Site 1 | UZH | 120 | 2 |
| *Klebsiella pneumoniae* | Aminoglycoside susceptible | 21 | Site 1 | UZH | 121 | 2 |
| *Klebsiella pneumoniae* | Aminoglycoside susceptible | 22 | Site 1 | UZH | 122 | 2 |
| *Klebsiella pneumoniae* | Aminoglycoside susceptible | 23 | Site 1 | UZH | 123 | 4 |
| *Klebsiella pneumoniae* | Aminoglycoside susceptible | 24 | Site 1 | UZH | 124 | 4 |
| *Klebsiella pneumoniae* | Aminoglycoside susceptible | 25 | Site 1 | UZH | 125 | 2 |
| *Klebsiella pneumoniae* | Aminoglycoside susceptible | 26 | Site 1 | UZH | 126 | 2 |
| *Klebsiella pneumoniae* | Aminoglycoside susceptible | 27 | Site 1 | UZH | 127 | 4 |
| *Klebsiella pneumoniae* | Aminoglycoside susceptible | 28 | Site 1 | UZH | 128 | 2 |
| *Klebsiella pneumoniae* | Aminoglycoside susceptible | 29 | Site 1 | UZH | 129 | 2 |
| *Klebsiella pneumoniae* | Aminoglycoside susceptible | 30 | Site 1 | UZH | 130 | 8 |
| *Klebsiella pneumoniae* | Aminoglycoside susceptible | 31 | Site 1 | UZH | 131 | 2 |
| *Klebsiella pneumoniae* | Aminoglycoside susceptible | 32 | Site 1 | UZH | 132 | 4 |
| *Klebsiella pneumoniae* | Aminoglycoside susceptible | 33 | Site 1 | UZH | 133 | 4 |
| *Klebsiella pneumoniae* | Aminoglycoside susceptible | 34 | Site 1 | UZH | 134 | 4 |
| *Klebsiella pneumoniae* | Aminoglycoside susceptible | 35 | Site 2 | RH | 1 | 2 |
| *Klebsiella pneumoniae* | Aminoglycoside susceptible | 36 | Site 2 | RH | 2 | 4 |
| *Klebsiella pneumoniae* | Aminoglycoside susceptible | 37 | Site 2 | RH | 3 | 4 |
| *Klebsiella pneumoniae* | Aminoglycoside susceptible | 38 | Site 2 | RH | 4 | 2 |
| *Klebsiella pneumoniae* | Aminoglycoside susceptible | 39 | Site 2 | RH | 6 | 2 |
| *Klebsiella pneumoniae* | Aminoglycoside susceptible | 40 | Site 2 | RH | 7 | 4 |
| *Klebsiella pneumoniae* | Aminoglycoside susceptible | 41 | Site 2 | RH | 8 | 4 |
| *Klebsiella pneumoniae* | Aminoglycoside susceptible | 42 | Site 2 | RH | 9 | 2 |
| *Klebsiella pneumoniae* | Aminoglycoside susceptible | 43 | Site 2 | RH | 10 | 4 |
| *Klebsiella pneumoniae* | Aminoglycoside susceptible | 44 | Site 2 | RH | 11 | 4 |
| *Klebsiella pneumoniae* | Aminoglycoside susceptible | 45 | Site 2 | RH | 12 | 4 |
| *Klebsiella pneumoniae* | Aminoglycoside susceptible | 46 | Site 2 | RH | 13 | 2 |
| *Klebsiella pneumoniae* | Aminoglycoside susceptible | 47 | Site 2 | RH | 14 | 2 |
| *Klebsiella pneumoniae* | Aminoglycoside susceptible | 48 | Site 2 | RH | 15 | 4 |
| *Klebsiella pneumoniae* | Aminoglycoside susceptible | 49 | Site 2 | RH | 16 | 2 |
| *Klebsiella pneumoniae* | Aminoglycoside susceptible | 50 | Site 2 | RH | 17 | 2 |
| *Klebsiella pneumoniae* | Aminoglycoside susceptible | 51 | Site 2 | RH | 18 | 4 |
| *Klebsiella pneumoniae* | Aminoglycoside susceptible | 52 | Site 2 | RH | 19 | 4 |
| *Klebsiella pneumoniae* | Aminoglycoside susceptible | 53 | Site 2 | RH | 20 | 4 |
| *Klebsiella pneumoniae* | Aminoglycoside susceptible | 54 | Site 2 | RH | 21 | 2 |
| *Klebsiella pneumoniae* | Aminoglycoside susceptible | 55 | Site 2 | RH | 22 | 4 |
| *Klebsiella pneumoniae* | Aminoglycoside susceptible | 56 | Site 2 | RH | 23 | 2 |
| *Klebsiella pneumoniae* | Aminoglycoside susceptible | 57 | Site 2 | RH | 24 | 8 |
| *Klebsiella pneumoniae* | Aminoglycoside susceptible | 58 | Site 2 | RH | 25 | 2 |
| *Klebsiella pneumoniae* | Aminoglycoside susceptible | 59 | Site 2 | RH | 26 | 4 |
| *Klebsiella pneumoniae* | Aminoglycoside susceptible | 60 | Site 2 | RH | 27 | 4 |
| *Klebsiella pneumoniae* | Aminoglycoside susceptible | 61 | Site 2 | RH | 28 | 4 |
| *Klebsiella pneumoniae* | Aminoglycoside susceptible | 62 | Site 2 | RH | 30 | 2 |
| *Klebsiella pneumoniae* | Aminoglycoside susceptible | 63 | Site 3 | UU | EN0092 | 1 |

**Table S2** (continued).

| Species | Phenoytpe | Isolate | Site | ID | Isolate ID | APR MIC (µg/mL) |
|---|---|---|---|---|---|---|
| *Klebsiella pneumoniae* | Aminoglycoside susceptible | 64 | Site 3 | UU | EN0094 | 1 |
| *Klebsiella pneumoniae* | Aminoglycoside susceptible | 65 | Site 3 | UU | EN0095 | 2 |
| *Klebsiella pneumoniae* | Aminoglycoside susceptible | 66 | Site 3 | UU | EN0096 | 2 |
| *Klebsiella pneumoniae* | Aminoglycoside susceptible | 67 | Site 3 | UU | EN0097 | 1 |
| *Klebsiella pneumoniae* | Aminoglycoside susceptible | 68 | Site 3 | UU | EN0098 | 2 |
| *Klebsiella pneumoniae* | Aminoglycoside susceptible | 69 | Site 3 | UU | EN0099 | 2 |
| *Klebsiella pneumoniae* | Aminoglycoside susceptible | 70 | Site 3 | UU | EN0100 | 0.5 |
| *Klebsiella pneumoniae* | Aminoglycoside susceptible | 71 | Site 3 | UU | EN0101 | 2 |
| *Klebsiella pneumoniae* | Aminoglycoside susceptible | 72 | Site 3 | UU | EN0102 | 2 |
| *Klebsiella pneumoniae* | Aminoglycoside susceptible | 73 | Site 3 | UU | EN0103 | 2 |
| *Klebsiella pneumoniae* | Aminoglycoside susceptible | 74 | Site 3 | UU | EN0104 | 1 |
| *Klebsiella pneumoniae* | Aminoglycoside susceptible | 75 | Site 3 | UU | EN0105 | 2 |
| *Klebsiella pneumoniae* | Aminoglycoside susceptible | 76 | Site 3 | UU | EN0106 | 2 |
| *Klebsiella pneumoniae* | Aminoglycoside susceptible | 77 | Site 3 | UU | EN0107 | 1 |
| *Klebsiella pneumoniae* | Aminoglycoside susceptible | 78 | Site 3 | UU | EN0109 | 2 |
| *Klebsiella pneumoniae* | Aminoglycoside susceptible | 79 | Site 3 | UU | EN0110 | 2 |
| *Klebsiella pneumoniae* | Aminoglycoside susceptible | 80 | Site 3 | UU | EN0112 | 2 |
| *Klebsiella pneumoniae* | Aminoglycoside susceptible | 81 | Site 3 | UU | EN0113 | 2 |
| *Klebsiella pneumoniae* | Aminoglycoside susceptible | 82 | Site 3 | UU | EN0115 | 2 |
| *Klebsiella pneumoniae* | Aminoglycoside susceptible | 83 | Site 3 | UU | EN0116 | 2 |
| *Klebsiella pneumoniae* | Aminoglycoside susceptible | 84 | Site 3 | UU | EN0197 | 2 |
| *Klebsiella pneumoniae* | Aminoglycoside susceptible | 85 | Site 3 | UU | EN0198 | 2 |
| *Klebsiella pneumoniae* | Aminoglycoside susceptible | 86 | Site 3 | UU | EN0199 | 1 |
| *Klebsiella pneumoniae* | Aminoglycoside susceptible | 87 | Site 3 | UU | EN0200 | 2 |
| *Klebsiella pneumoniae* | Aminoglycoside susceptible | 88 | Site 3 | UU | EN0201 | 1 |
| *Klebsiella pneumoniae* | Aminoglycoside susceptible | 89 | Site 3 | UU | EN0202 | 2 |
| *Klebsiella pneumoniae* | Aminoglycoside susceptible | 90 | Site 3 | UU | EN0204 | 1 |
| *Klebsiella pneumoniae* | Aminoglycoside susceptible | 91 | Site 3 | UU | EN0205 | 2 |
| *Klebsiella pneumoniae* | Aminoglycoside susceptible | 92 | Site 3 | UU | EN0206 | 1 |
| *Klebsiella pneumoniae* | Aminoglycoside susceptible | 93 | Site 3 | UU | EN0207 | 2 |
| *Klebsiella pneumoniae* | Aminoglycoside susceptible | 94 | Site 3 | UU | EN0208 | 1 |
| *Klebsiella pneumoniae* | Aminoglycoside susceptible | 95 | Site 3 | UU | EN0209 | 4 |
| *Klebsiella pneumoniae* | Aminoglycoside susceptible | 96 | Site 3 | UU | EN0210 | 1 |
| *Klebsiella pneumoniae* | Aminoglycoside susceptible | 97 | Site 3 | UU | EN0211 | 1 |
| *Klebsiella pneumoniae* | Aminoglycoside susceptible | 98 | Site 3 | UU | EN0212 | 1 |
| *Klebsiella pneumoniae* | Aminoglycoside susceptible | 99 | Site 3 | UU | EN0213 | 1 |
| *Klebsiella pneumoniae* | Aminoglycoside susceptible | 100 | Site 3 | UU | EN0214 | 1 |
| *Klebsiella pneumoniae* | Aminoglycoside susceptible | 101 | Site 3 | UU | EN0253 | 2 |
| *Klebsiella pneumoniae* | Aminoglycoside susceptible | 102 | Site 3 | UU | EN0254 | 1 |
| *Klebsiella pneumoniae* | Aminoglycoside susceptible | 103 | Site 3 | UU | EN0256 | 1 |
| *Klebsiella pneumoniae* | Aminoglycoside susceptible | 104 | Site 3 | UU | EN0257 | 1 |
| *Klebsiella pneumoniae* | Aminoglycoside susceptible | 105 | Site 3 | UU | EN0258 | 2 |
| *Klebsiella pneumoniae* | Aminoglycoside susceptible | 106 | Site 4 | IHMA | 2181270 | 2 |
| *Klebsiella pneumoniae* | Aminoglycoside susceptible | 107 | Site 4 | IHMA | 2315139 | 2 |
| *Klebsiella pneumoniae* | Aminoglycoside susceptible | 108 | Site 4 | IHMA | 2008168 | 2 |
| *Klebsiella pneumoniae* | Aminoglycoside susceptible | 109 | Site 4 | IHMA | 1987016 | 2 |
| *Klebsiella pneumoniae* | Aminoglycoside susceptible | 110 | Site 4 | IHMA | 1995157 | 4 |
| *Klebsiella pneumoniae* | Aminoglycoside susceptible | 111 | Site 4 | IHMA | 1989403 | 2 |
| *Klebsiella pneumoniae* | Aminoglycoside susceptible | 112 | Site 4 | IHMA | 2015467 | 4 |
| *Klebsiella pneumoniae* | Aminoglycoside susceptible | 113 | Site 4 | IHMA | 1971691 | 2 |
| *Klebsiella pneumoniae* | Aminoglycoside susceptible | 114 | Site 4 | IHMA | 1973885 | 2 |
| *Klebsiella pneumoniae* | Aminoglycoside susceptible | 115 | Site 4 | IHMA | 1957286 | 2 |
| *Klebsiella pneumoniae* | Aminoglycoside susceptible | 116 | Site 4 | IHMA | 2007378 | 2 |
| *Klebsiella pneumoniae* | Aminoglycoside susceptible | 117 | Site 4 | IHMA | 1970636 | 4 |
| *Klebsiella pneumoniae* | Aminoglycoside susceptible | 118 | Site 4 | IHMA | 1963657 | 2 |
| *Klebsiella pneumoniae* | Aminoglycoside susceptible | 119 | Site 4 | IHMA | 1993652 | 4 |
| *Klebsiella pneumoniae* | Aminoglycoside susceptible | 120 | Site 4 | IHMA | 2058859 | 2 |
| *Klebsiella pneumoniae* | Aminoglycoside susceptible | 121 | Site 4 | IHMA | 2025423 | 2 |
| *Klebsiella pneumoniae* | Aminoglycoside susceptible | 122 | Site 4 | IHMA | 2025196 | 2 |
| *Klebsiella pneumoniae* | Aminoglycoside susceptible | 123 | Site 4 | IHMA | 2089546 | 2 |
| *Klebsiella pneumoniae* | Aminoglycoside susceptible | 124 | Site 4 | IHMA | 1986505 | 1 |
| *Klebsiella pneumoniae* | Aminoglycoside susceptible | 125 | Site 4 | IHMA | 2227489 | 2 |

**Table S3.** Frequency of mutational resistance to apramycin within 48 hours of incubation.

| Species /Strain | Resistance gene annotations include | Drug | MIC (µg/mL) | Frequency of resistance | |
|---|---|---|---|---|---|
| | | | | 4 × MIC | 8 × MIC |
| ***E. coli*** | | | | | |
| ATCC 25922 | | APR | 8 | <4.5E-10 | <4.5E-10 |
| | | GEN | 1 | 2.2E-09 | <4.75E-10 |
| EN0591 | CTX, *rmtB, aac(3)-II, aph(3')-II, aadA* | APR | 8 | 1.0E-09 | <4.9E-11 |
| EN1107 | NDM5, *rmtB, aac(3)-II, aac(6')-Ib, aadA* | APR | 4 | 5.3E-10 | <1.3E-10 |
| EN1108 | NDM5, *rmtB, aac(3)-II, aac(6')-Ib, aadA* | APR | 4 | 2.9E-09 | <2.1E-10 |
| EN1109 | NDM1, *rmtC, aac(6')-Ib, aadA* | APR | 8 | 1.6E-09 | <1.6E-10 |
| EN1110 | NDM5, *rmtB, aac(3)-II, aadA* | APR | 4 | 2.7E-08 | 4.8E-10 |
| EN1111 | NDM1, *armA, aac(3)-II, aac(6')-Ib, aadA* | APR | 4 | 8.4E-08 | <4.5E-10 |
| EN1112 | NDM7, *rmtB, aac(6')-Ib, aadA* | APR | 4 | 3.5E-09 | <4.3E-10 |
| ***K. pneumoniae*** | | | | | |
| ATCC 13883 | | APR | 4 | <3.8E-10 | <3.8E-10 |
| | | GEN | 0.125 | 4.1E-08 | <7.9E-10 |
| EN0593 | NDM1, *rmtB, aac(3)-II, aac(6')-Ib, aadA* | APR | 4 | <4.4E-10 | <4.4E-10 |
| EN1113 | NDM1, *rmtC, aac(6')-Ib* | APR | 4 | 9.5E-10 | < 4.8E-10 |
| EN1114 | NDM1, *rmtC, aac(6')-Ib, aph(3')-I* | APR | 4 | 3.1E-09 | < 6.3E-10 |
| EN1115 | NDM1, *rmtF, aac(6')-Ib, aph(3')-I, aadA* | APR | 2 | 2.6E-10 | < 2.6E-10 |
| EN1116 | NDM1, *rmtC, rmtF, aac(6')-Ib, aph(3')-I, aadA* | APR | 4 | < 1.7E-10 | < 1.7E-10 |
| EN1117 | NDM1, *rmtC, aac(6')-Ib, aph(3')-I* | APR | 2 | 5.0E-10 | < 2.5E-10 |
| EN1118 | NDM1, *armA, rmtF, aac(3)-II, aac(6')-Ib, aph(3')-I, aph(3')-VI, ant(3'')-I, aadA* | APR | 4 | 3.4E-10 | < 3.4E-10 |
| ***E. cloacae*** | | | | | |
| EN0594 | *rmtC, aac(3)-II, aac(6')-Ib, aph(6)-I* | APR | 4 | <4.7E-10 | <4.7E-10 |

**Table S4**. Data on 20 CPE strains (8 *E.coli* and 12 *K. pneumoniae* isolates) selected according to WGS to represent different genotypes for carbapenemase-production.

| Isolate | Carbapenemase genes | Other β-lactamase genes | Aminoglycoside resistance genes | MIC (µg/mL) | Disk diffusion interpretation | | |
|---|---|---|---|---|---|---|---|
| | | | | APR | GEN | TOB | AMK |
| EC WT [1] | None | None | None | 4 | S | S | S |
| EC1 | *bla*VIM-1 | *bla*TEM-1B, *bla*OXA-1, *bla*CTX-M-15 | *aac(3)-IIa* | 8 | R | R | R |
| EC2 | *bla*NDM-1 | *bla*OXA-1, *bla*CMY-6-like, *bla*DHA-1-like | *rmtC, aph(6)-Id, aph(3'')-Ib*-like | 8 | R | R | R |
| EC3 | *bla*NDM-5 | *bla*OXA-1, *bla*CTX-M-15, *bla*CMY-42 | *aac(3)-IIa*-like, *aph(6)-Id, aph(3'')-Ib*-like | 4 - 8 | R | R | S |
| EC4 | *bla*NDM-5, OXA-181 | *bla*TEM-1B, *bla*OXA-1, *bla*CTX-M-15, *bla*CMY-2 | *aac(3)-IId*-like, *aadA2*-like, *aph(6)-Id* | 4 | R | R | S |
| EC5 | *bla*NDM-7 | *bla*TEM-1b, *bla*-CTX-M-27 | *aac(3)-IId*-like, *aph(6)-Id*-like, *aph(3'')-Ib*-like | 8 | R | R | S |
| EC6 | *bla*OXA-244 | *bla*TEM-1B, *bla*CTX-M-14b | *aph(3')-Ia, aph(6)-Id*-like, *aph(3'')-Ib*-like | 8 - 16 | S | S | S |
| EC7 | *bla*OXA-48 | *bla*TEM-1B-like | *aadA5* | 4 | S | S | S |
| EC8 | *bla*VIM-4 | *bla*OXA-1, *bla*CTX-M-15, *bla*CTX-M-9 | *aac(3)-IIa*-like, *aac(6')-Il* | 8 | R | R | R |
| KP WT [2] | None | None | None | 2 | S | S | S |
| KP1 | *bla*KPC-2 | *bla*TEM-1A-like, *bla*OXA-9-like | *aph(3')-Ia* | 2 | S | R | R |
| KP2 | *bla*KPC-3 | *bla*TEM-1B, *bla*SHV-11, *bla*OXA-1, *bla*OXA-9-like | *aac(3)-IId*-like, *aph(6)-Id* | 2 | R | R | R |
| KP3 | *bla*KPC-3 | *bla*SHV-11-like | *aac(6')-Ib, aac(3)-IVa*-like, *aadA2*-like | > 64 | R | R | R |

[1] EC WT, *E. coli* ATCC 25922

[2] KP WT, *K. pneumoniae* ATCC 13883

**Table S4** (continued).

| Isolate | Carbapenemase genes | Other β-lactamase genes | Aminoglycoside resistance genes | MIC (µg/mL) | Disk diffusion interpretation | | |
|---|---|---|---|---|---|---|---|
| | | | | **APR** | **GEN** | **TOB** | **AMK** |
| KP4 | *bla*KPC-3 | *bla*SHV-1 | *armA* | 2 | R | R | R |
| KP5 | *bla*KPC-3 | *bla*TEM-1B-like, *bla*SHV-28, *bla*OXA-1, *bla*OXA-9-like, *bla*CTX-M-15 | *aac(3)-IIa*-like, *aph(6)-Id, aph(3'')-Ib* | 4 | R | R | S |
| KP6 | *bla*NDM-1 | *bla*TEM-1B, *bla*SHV-11-like, *bla*OXA-1, *bla*CTX-M-15, *bla*CMY-6, *bla*DHA-1-like | *rmtC, aac(3)-IIa*-like, *aph(6)-Id, aph(3'')-Ib* | 2 | R | R | R |
| KP7 | *bla*NDM-1, *bla*OXA-48 | *bla*TEM-1B, *bla*SHV-12, *bla*OXA-1, *bla*CTX-M-15 | *rmtC, aac(3)-IIa*-like, *aadA5* | 2 | R | R | R |
| KP8 | *bla*NDM-5 | *bla*TEM-1B, *bla*SHV-1, *bla*OXA-1, *bla*CTX-M-15 | *aph(6)-Id, aph(3'')-Ib* | 2 | S | R | S |
| KP9 | *bla*NDM-7, *bla*OXA-181 | *bla*TEM-1B, *bla*SHV-11-like, *bla*CTX-M-15, *bla*CMY-6 | *rmtC, rmtF aadA2*-like, *aph(6)-Id, aph(3'')-Ib* | 2 | R | R | R |
| KP10 | *bla*OXA-232 | *bla*TEM-1B, *bla*SHV-1, *bla*CTX-M-15 | *rmtF*-like *aadA2*-like | 4 | R | R | R |
| KP11 | *bla*OXA-436 | *bla*TEM-1B, *bla*SHV-1-like, *bla*OXA-10 | *aac(6')-IIc*-like, *aadA2*-like, *aph(6)-Id, aph(3'')-Ib*-like | 2 | R | R | S |
| KP12 | *bla*IMP-1 | *bla*SHV-26 | *aac(6')-Im* | 8 | R | R | R |

[1] EC WT, *E. coli* ATCC 25922

[2] KP WT, *K. pneumoniae* ATCC 13883

**Table S5**. Characterization of apramycin-resistant colonies obtained in the FoR studies

| Species | Strain/Clone | Doubling time (min) | Relative fitness | Sequencing results and interpretation | Reference |
|---|---|---|---|---|---|
| *E. coli* | EN591 (WT) | 31.4 | 1.00 | Wild type | |
| *E. coli* | CH7440 | 44.9 | 0.70 | IS insertion 139 bp upstream of cydA. Small colony variant. | [1, 2] |
| *E. coli* | CH7441 | 33.4 | 0.94 | ND | |
| *E. coli* | CH7442 | 52.9 | 0.59 | ubiD W415* Small colony variant. | [3] |
| *E. coli* | CH7443 | 37.5 | 0.84 | cydA M166::IS Small colony variant. | [1, 2] |

*ND, not determined*
